# A transcriptomic and spatial map of serotonin autoreceptor expression in *Drosophila*

**DOI:** 10.64898/2026.08.27.747575

**Authors:** Luke Brewer, Ye Li, Juri Lim, Solmari Solano, Emma Walker, Megan Watts, Jessie K. Rhodes, Dawen Cai, Douglas H. Roossien

## Abstract

Serotonin is an evolutionarily ancient neurotransmitter that modulates an array of behaviors such as mood, sleep, and appetite across species. Serotonin acts primarily by binding to serotonin receptors, which are expressed in post-synaptic neurons (heteroreceptors) and serotonergic neurons themselves (autoreceptors). Serotonin autoreceptors modulate serotonergic tone, the foundational principles of which have been excellently demonstrated in vertebrate and invertebrate models. However, many aspects of the mechanisms and contexts in which this modulation occurs are still unclear. *Drosophila melanogaster* is a powerful model organism that can provide unique insights into autoreceptor function by the ability to perform precise spatial and temporal genetic manipulation with structural and functional readouts / behaviors of serotonin systems. However, a systematic characterization of serotonin autoreceptor expression in *Drosophila* has not been conducted. Here we use single-cell sequencing and genetic labeling to show that all five serotonin receptors are expressed in serotonergic neurons and map their expression at both the larval and adult stages of development. This is the first evidence of 5-HT2A and 5-HT7 expression in serotonergic neurons in any organism. Moreover, the unique combinations of autoreceptor expression in specific neuronal clusters will aid in the development of novel hypotheses for autoreceptor function, and demonstrates the utility of *Drosophila* as a model organism to study the function of serotonin autoreceptors.

**Significance Statement:** This paper is a valuable addition to the neuroscientific community because it: 1) is the first to discover all serotonin receptors are autoreceptors in *Drosophila*; 2) is the first to identify 5-HT2A and 5-HT7 autoreceptor expression in any organism; 3) demonstrates putative unique combinatorial autoreceptor expression in certain serotonergic neurons; 4) provides spatial information as to which serotonergic neurons express autoreceptors, which will support the development of future testable hypotheses for autoreceptor function in *Drosophila*.

## Introduction

Serotonin receptors are categorized into 7 major families (5-HT1-7) based on structure and function. 5-HT3 receptors are ligand-gated ion channels, while the rest are metabotropic GPCRs. Although many serotonin receptors have emerged throughout evolution, members of the 5-HT1, 5-HT2, and 5-HT7 families are highly conserved in both structure and function between vertebrates and invertebrates. For example, the *Drosophila* genome contains five serotonin receptors: 5-HT1A, 5-HT1B, 5-HT2A, 5-HT2B, and 5-HT7, all named based on their closest vertebrate homologs. All of these metabotropic serotonin receptors couple to signaling pathways that activate second messenger cascades to modulate ion channels and membrane polarization. Serotonin receptors can be expressed in both post-synaptic non-serotonergic neurons (referred to as heteroreceptors) and on serotonergic neurons themselves (referred to as autoreceptors).

The two subpopulations make unique contributions to cell signaling, electrophysiology, and behavior. For example, 5-HT1A heteroreceptors modulate a variety of behaviors based on their expression in specific brain regions, whereas 5-HT1A autoreceptors suppress 5-HT release throughout the brain via autoinhibition (Albert & Vahid-Ansari, 2019; la Cour et al., 2006; Sharp & Barnes, 2020).

Differences in signaling underlie unique physiological and behavioral roles for serotonin autoreceptors compared to their heteroreceptor counterparts (Garcia-Garcia et al., 2014). In vertebrates, selective 5-HT1A autoreceptor activation leads to anxiolytic effects (Richardson-Jones et al., 2010, 2011). Disrupting 5-HT1A autoreceptors selectively during adulthood creates an anxiogenic effect in animals treated with antidepressants (Turcotte-Cardin et al., 2019), and elevated 5-HT1A autoreceptor expression underlies depression in humans and mice (Albert et al., 2014; Garcia-Garcia et al., 2014). Similar findings have been reported for 5-HT1B as well: 5- HT1B autoreceptor knockout mice display antidepressant phenotypes whereas 5-HT1B heteroreceptor knockouts do not (Nautiyal et al., 2016). The cellular mechanisms of autoregulation of serotonin (5-Hydroxytryptamine or 5-HT) release has been studied extensively in cultured leech neurons (Trueta & Cercós, 2026), though those cannot be directly correlated to changes in circuit function or behavior. Establishing *Drosophila* as a model organism to study serotonin autoreceptors can help bridge this gap by providing a system with precise genetic manipulations together with cellular and functional readouts.

In vertebrates, 5-HT1A and 5-HT1B are the most prevalent autoreceptors, with evidence that low amounts of 5-HT2B are expressed in the raphe as well (Sharp & Barnes, 2020). There have been reports in *Drosophila* that 5-HT1A autoreceptors can be found in adult brains (Alekseyenko et al., 2019; Sampson et al., 2020; Yuan et al., 2005), are minimally expressed in larvae (Huser et al., 2017), and in subsets of cultured embryonic *Drosophila* serotonergic neurons (Long et al., 2023).

Together this suggests 5-HT1A autoreceptor expression is persistent throughout development, though whether the remaining receptors are is unknown. Moreover, these previous reports identifying serotonin autoreceptors in *Drosophila* relied on genetic labeling strategies, which may not be sensitive enough to detect receptors expressed at low levels or transiently. Here, analysis of single-cell sequencing data showed that all five serotonin receptors are expressed in serotonergic neurons at both the 3rd instar larval stage and adult stage. By crossing LexA drivers for each serotonin receptor to a fluorescent reporter and co-staining with 5-HT antibodies, we created a spatial map of serotonin receptor-expressing serotonergic neurons at these stages. As *Drosophila* is a highly valuable and genetically tractable model organism, understanding the expression of serotonin receptors throughout the serotonergic system will aid in the development of future hypotheses relating to autoreceptor function and behavior.

## Methods

### Fly husbandry

Flies were raised on standard Bloomington Formula NutriFly *Drosophila* media (Genesee #66- 121) at 25°C with 12:12 hr light:dark cycling. Genetic labeling for molecular and spatial mapping was accomplished by crossing 5-HTR-LexA driver flies to UAS-mCD8::RFP, LexAop-mCD8::GFP reporter flies (Pfeiffer et al., 2010). Genetic labeling for cell sorting and single-cell sequencing was achieved by crossing the TRH-Gal4 driver line (Alekseyenko et al., 2010) to the nucleus-localized mNeonGreen reporter line (Michki et al., 2021). All flies were obtained from the Bloomington *Drosophila* Stock Center. See Table 1 for a complete list of stock genotypes and numbers used in this study.

**Table 1:** Fly stocks used in this study.

| Line | Genotype | Stock #<br>(BDSC) | RRID |
| --- | --- | --- | --- |
| GFP reporter | y[1] w[*] P{y[+t7.7] w[+mC]=10XUAS-IVS-mCD8::RFP}attP18 P{y[+t7.7] w[+mC]=13XLexAop2-mCD8::GFP}su(Hw)attP8 | 32229 | RRID:BDSC_32229 |
| 5-HT1A-LexA | w[*]; Tl{2A-lexA::GAD}5-HT1A[2A-lexA]/CyO | 84350 | RRID:BDSC_84350 |
| 5-HT1B-LexA | Tl{2A-lexA::GAD}5-HT1B[2A-AD.lexA] | 84351 | RRID:BDSC_84351 |
| 5-HT2A-LexA | Tl{2A-lexA::GAD}5-HT2A[2A-AEC.lexA] | 84352 | RRID:BDSC_84352 |
| 5-HT2B-LexA | w[*]; Tl{2A-lexA::p65}5-HT2B[2A-D.lexA] | 84353 | RRID:BDSC_84353 |
| 5-HT7-LexA | Tl{2A-lexA::GAD}5-HT7[2A-lexA] | 84354 | RRID:BDSC_84354 |
| nuc-mNG | y[1] w[*]; P{y[+t7.7] w[+mC]=UAS-hH2B.2xmNG}attP2 | 99280 | RRID:BDSC_99280 |
| TRH-Gal4 | w[1118]; P{w[+mC]=Trhn-GAL4.long}2 | 38388 | RRID:BDSC_38388 |

### Single-cell sequencing

Serotonergic neurons were labeled in the progeny larvae from crosses of transgenic lines TRH- Gal4 (Alekseyenko et al., 2010) and UAS-H2B-2xNeonGreen (Michki et al., 2021). Specificity of the labeling was confirmed by cell body positions through imaging (**Fig 1A**). A total of 53 late 3rd instar larval brains were dissected in 1 hour and quickly proceeded to brain dissociation. The dissociation was effective based on imaging quantification of the cell population (statistics in **Table 2**), where 3960, or 93.4% of the NG+ cells, were retained (53x80=4240 total NG+ cells as 100% in estimation). The cells were then selected by FACS, where 1360 NG+ cells (32.08% of estimated total) were collected. After a quick wash and gentle pelleting down, a final 1140 of NG+ cells (26.89% of estimated total) were collected and proceeded to the 10X Chromium v3 pipeline to generate the single-cell RNA library. The library was sequenced on the NovaSeq 6000 platform, and yielded a total of 600M paired-end reads with 28bp for the cell barcode and UMI and 115bp for cDNA inserts.

**Figure 1:**
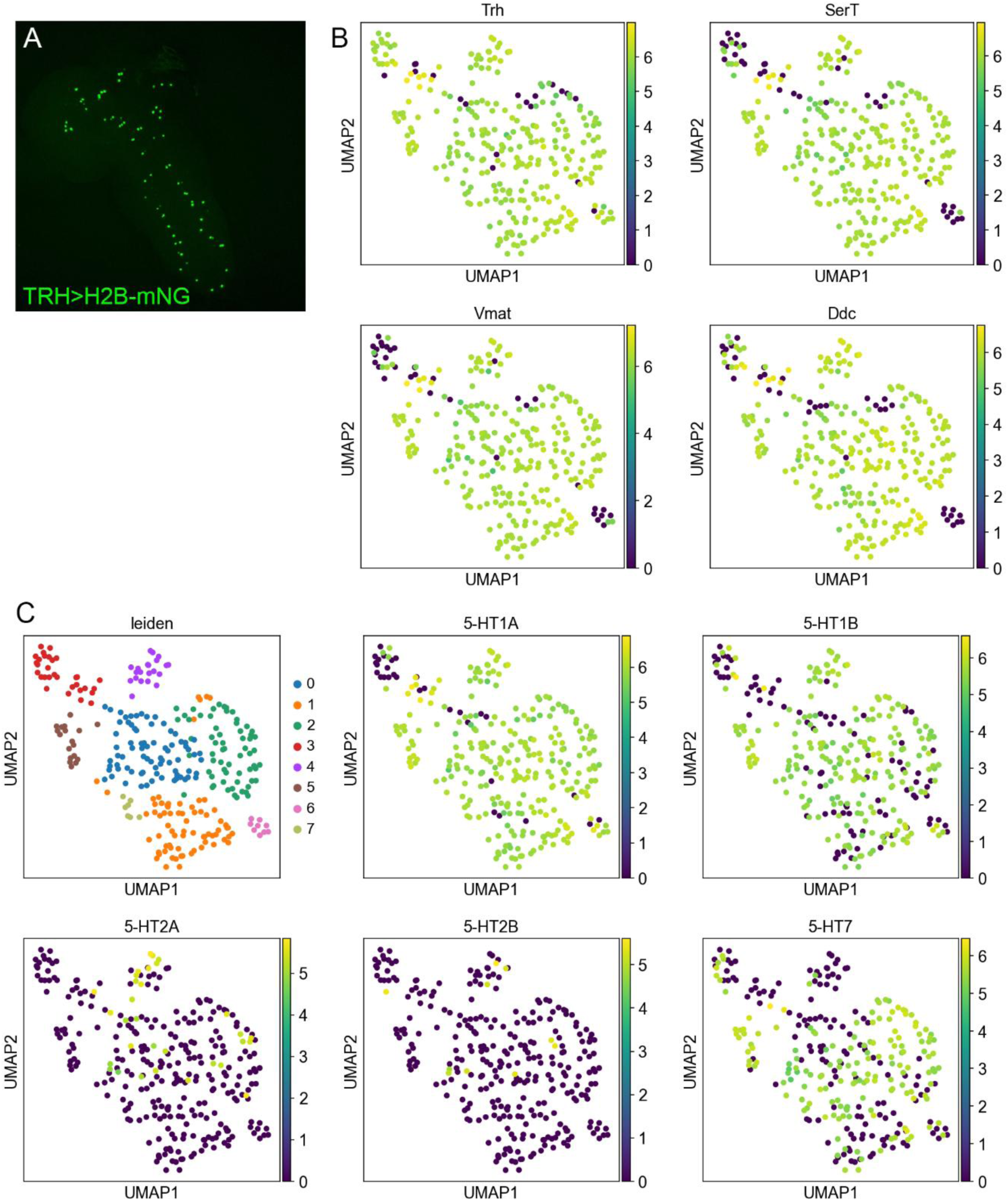
Single-cell sequencing of larval serotonergic neurons shows expression of all five serotonin receptors. (**A**) Larval serotonergic neurons labeled by nucleus-localized mNeonGreen. 3rd instar larva brain from the cross between TRH-Gal4 and UAS-H2B- 2xmNeonGreen. Brain is anterior-posterior organized from top-left to bottom-right. (**B**) Confirmation of classic marker genes related to serotonergic cells. Trh (Tryptophan hydroxylase) and Ddc (Dopa decarboxylase) are in the serotonin synthesis pathway. SerT (Serotonin transporter) and Vmat (Vesicular monoamine transporter) are related to serotonin transport. (**C**) Leiden cluster identity of the sequenced larval serotonergic neurons, as well as the expression distribution patterns of all five serotonin receptors, are displayed with a UMAP embedding in 2D.

**Table 2:** Cell counts in larval dissociation and sorting steps before sequencing.

| Step | Cell Count | % of previous step | % of first step |
| --- | --- | --- | --- |
| Dissection | 4240 (estimate) | - | 100% |
| Dissociation | 3960 | 93.40% | 93.40% |
| FACS Sorting | 1812 | 45.76% | 42.74% |
| Count after FACS | 1360 | 75.06% | 32.08% |
| Centrifuge & transfer | 1140 | 83.82% | 26.89% |

### Sequencing analysis

Reads were mapped using STAR-solo (Dobin et al., 2013) to the *Drosophila* genome assembly provided by ENSEMBL, build BDGP6 (2014-07). The downstream scRNA-seq analysis was performed using scanpy (Wolf et al., 2018). With preliminary filtering, 299 cells (7.05% of the estimated total) were present in the dataset with, at the median, >16,000 UMI counts and 3824 genes detected per cell. Based on the estimation of 80 total serotonergic neurons in the larval nervous system, and an assumption that our cell selection method was unbiased, our data represented a 3.73x coverage of targeted larval serotonergic neurons. For data processing, cells were filtered by requiring at least 200 unique genes/cell, and genes were filtered by requiring at least 2 cells to express it at greater than 1 UMI/cell. UMI counts were normalized to a total sum of 1e6 counts/cell (conversion to counts-per-million/CPM) and subsequently log-transformed by calculating ln(1+CPM) for each gene for each cell. The top 2000 highly variable genes were identified using the cell-ranger method and these genes were used to perform a principal component analysis (PCA, n = 50 PCs). The data then proceeded to neighborhood identification (k = 20), and finally a UMAP projection (2D). Clusters were identified using the Leiden algorithm (Traag et al., 2019) with clustering resolution at 0.5. Marker genes were identified using logistic regression analysis, implemented in scanpy.

### Immunofluorescence

Brains from third instar larvae or adults were dissected in PBS, fixed in 4% paraformaldehyde for 20 min at room temperature, then washed three times in PBS. Brains were blocked in Thermo StartingBlock-PBS with 1.0% Triton X-100 for 1-2 hr at room temperature followed by three washes in PBS with 0.3% Triton X-100 for 15 min each. Primary rabbit anti-serotonin antibody (ImmunoStar #20080; RRID AB_572263) and donkey anti-rabbit IgG TRITC secondary (Jackson ImmunoResearch #711-025-152; RRID AB_2340588) were prepared at 1:500 dilution in PBS with 0.3% Triton X-100. Samples were incubated in primary for 2-3 days at 4°C, and secondary for 1- 2 days at 4°C, with three 30 min washes in PBS with 0.3% Triton X-100 after both antibody incubations. All steps were performed with gentle rocking. Brains were mounted in Vectashield PLUS antifade medium (Vector Laboratories H-1900).

### Microscopy and image scoring

Larval samples from 5-HT1A>GFP and 5-HT2A>GFP F1s were imaged on a Nikon C2 confocal microscope, using 488 and 561 laser lines for GFP and TRITC excitation, respectively, and collected with a 20x NA 0.75 objective. The remaining brains were imaged on a Zeiss Apotome Axio Imager.M2 with a 20x NA 0.8 objective, X-cite 120LED Lamp, and Zeiss 807 mono camera. GFP was excited at 450 - 490 nm with emission collected at 500 - 550 nm; TRITC was excited at 530 - 560 nm with emission collected at 575 - 640 nm. On both systems, image stacks were collected with 2 µm Z-steps and 225 - 300 nm pixel size in x,y. At least four brains from each genotype and stage were independently scored using FIJI for double-positive signal in both GFP and TRITC channels by three individuals. A cluster was considered a “positive hit” with a consensus of at least two individuals in at least 75% of the brains. Anatomical reference maps were created using previously published descriptions for larvae (Huser et al., 2012) and adult brains (Pooryasin & Fiala, 2015).

## Results

We first wanted to ask whether serotonin receptors are expressed in serotonergic neurons using single-cell sequencing, as it is more sensitive than previously published genetic labeling approaches. To enrich a cell population for serotonergic neurons, we sorted fluorescently tagged serotonergic neurons and performed single-cell sequencing. Neurons were tagged by driving expression of UAS-H2B::mNeonGreen with the TRH-Gal4 driver, which expresses selectively in serotonergic neurons (Alekseyenko et al., 2010). A total of 53 dissected brains were quickly processed into single-cell suspensions and sorted by fluorescence using a Sony MA800S followed by 10X Chromium v3single-cell sequencing. Out of an expected cell count of 4,240 neurons (estimated from 53 larval brains), we successfully sorted and preserved 1140 serotonergic neurons, a 26.89% yield, for single-cell sequencing (**Table 2**), which achieved a deep read-depth dataset containing 299 cells after necessary filtering. We found 93.6% expressed tryptophan hydroxylase (TRH) and 88.3% expressed serotonin transporter (SerT), suggesting our dataset was highly enriched for serotonergic neurons (**Fig 1B**). Next, we asked whether serotonin receptors were expressed in these cells and found each of the five serotonin receptors. 5-HT1A, 5-HT1B, and 5-HT7 were expressed in the majority of serotonergic neurons, whereas 5-HT2A and 5-HT2B were rare (**Fig 1D-H**). Interestingly, none of the receptors correlated with any of the 7 clusters, suggesting that serotonin autoreceptor expression may not be driven by subset-specific genetic programs. Nonetheless, these data suggest all five serotonin receptors are expressed as autoreceptors in the third instar larval stage.

Next, we sought to determine which serotonergic neurons in the larval brain express serotonin autoreceptors. The *Drosophila* larval brain contains a stereotypical serotonergic system composed of ∼84 neurons (Huser et al., 2012; Vallés & White, 1988) each with a known anatomical location. This allows us to make systematic comparisons of expression patterns across multiple brains. First we attempted to drive UAS-GFP using 5-HTR-Gal4 drivers for each serotonin receptor. When we stained for serotonergic neurons using ɑ5HT antibodies, we found very few double positive cells with the 5-HT1B-Gal4 and 5-HT2B-Gal4 drivers, and those we did find had very weak fluorescence (data not shown). Based on our single-cell sequencing data it suggested the drivers may not be strong enough to accurately label weakly expressing cells. We therefore used a panel of 5-HTR-LexA drivers to genetically label receptor-expressing neurons with LexAop-GFP. Neurons double-positive for GFP and ɑ5HT (**Fig 2B**) were scored from at least four brains for each genotype based on anatomical position. We considered neurons that scored as double-positive in at least 75% of brains to be an autoreceptor-expressing serotonergic neuron. We found 5-HT1A expressed in serotonergic neurons in regions of the optic lobes: LP (**Fig 2B**), SP1 (**Fig 2C**), SP2 (**Fig 2D**), and IP (**Fig 2E**); whereas no serotonergic neurons in the ventral nerve cord (VNC) appeared to express 5-HT1A. 5-HT1B was expressed in the SP2 (**Fig 3B**) and IP (**Fig 3C**) clusters, but also in SE1 and SE3 in the VNC (**Fig 3D-E**). 5-HT2B was expressed in SP1 (**Fig 4B**), IP (**Fig 4C**), and LP1 (**Fig 4D**). 5-HT7 was expressed in SP1 (**Fig 5B**), SP2 (**Fig 5C**) and LP1 (**Fig 5D**) of the optic lobes, but also in A2-8/9 in the VNC (**Fig 5E-F**). Surprisingly, we did not find any double-positive neurons from the 5-HT2A line. A summary of these results can be found in Table 3.

**Figure 2:**
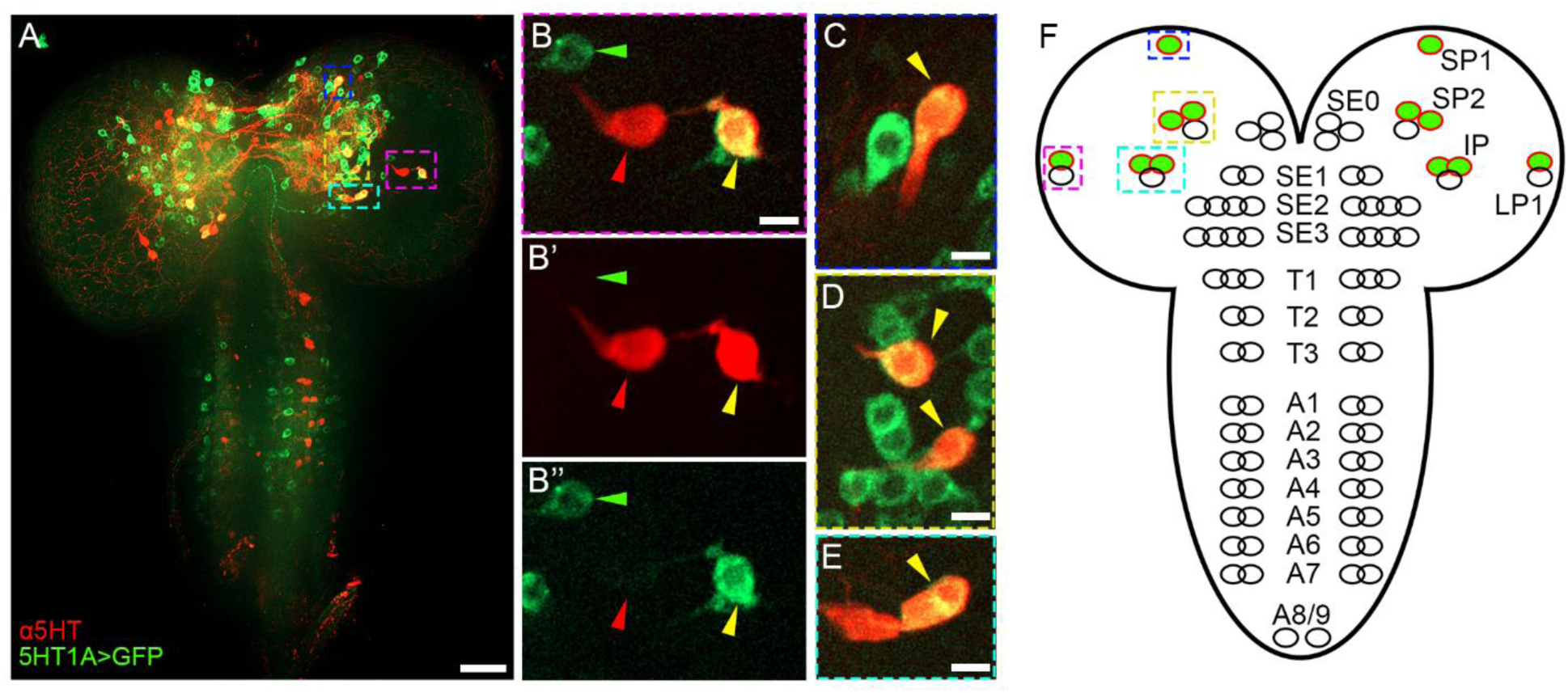
Spatial map of serotonin autoreceptor 5-HT1A expression in third instar brains by genetic labeling. (**A-E**) Images from 5-HT1A-LexA; LexAop-GFP brains stained with ɑ5HT antibodies. (**A**) Maximum projection of the Z-stack shows the anatomical layout of the serotonergic system (red) together with neurons that express 5-HT1A (green). (**B-B’’**) Magnified view of the LP1 cluster. (B) Merge of the two channels shows an example of a non-serotonergic neuron expressing 5-HT1A (green arrowhead), a serotonergic neuron that does not express 5- HT1A (red arrowhead), and a serotonergic neuron that does express 5-HT1A (yellow arrowhead), as evidenced by colabeling of GFP and ɑ5-HT antibody. (**C**) SP1 serotonergic neurons expressing 5-HT1A. (**D**) SP2 serotonergic neurons expressing 5-HT1A. (**E**) IP neurons expressing 5-HT1A. (**F**) Schematic of a larval brain with approximated serotonergic clusters. Neurons colorized red and green indicate identified serotonergic neurons expressing 5-HT1A in at least 75% of brains scored. Incompletely colorized clusters indicate some but not all serotonergic neurons express 5- HT1A in that cluster, but do not necessarily represent the specific anatomical position within that cluster. Colored boxes in (A-F) identify the same clusters across panels. Scale bar in (A) = 25 µm; scale bar in (B-E) = 5 µm.

**Figure 3:**
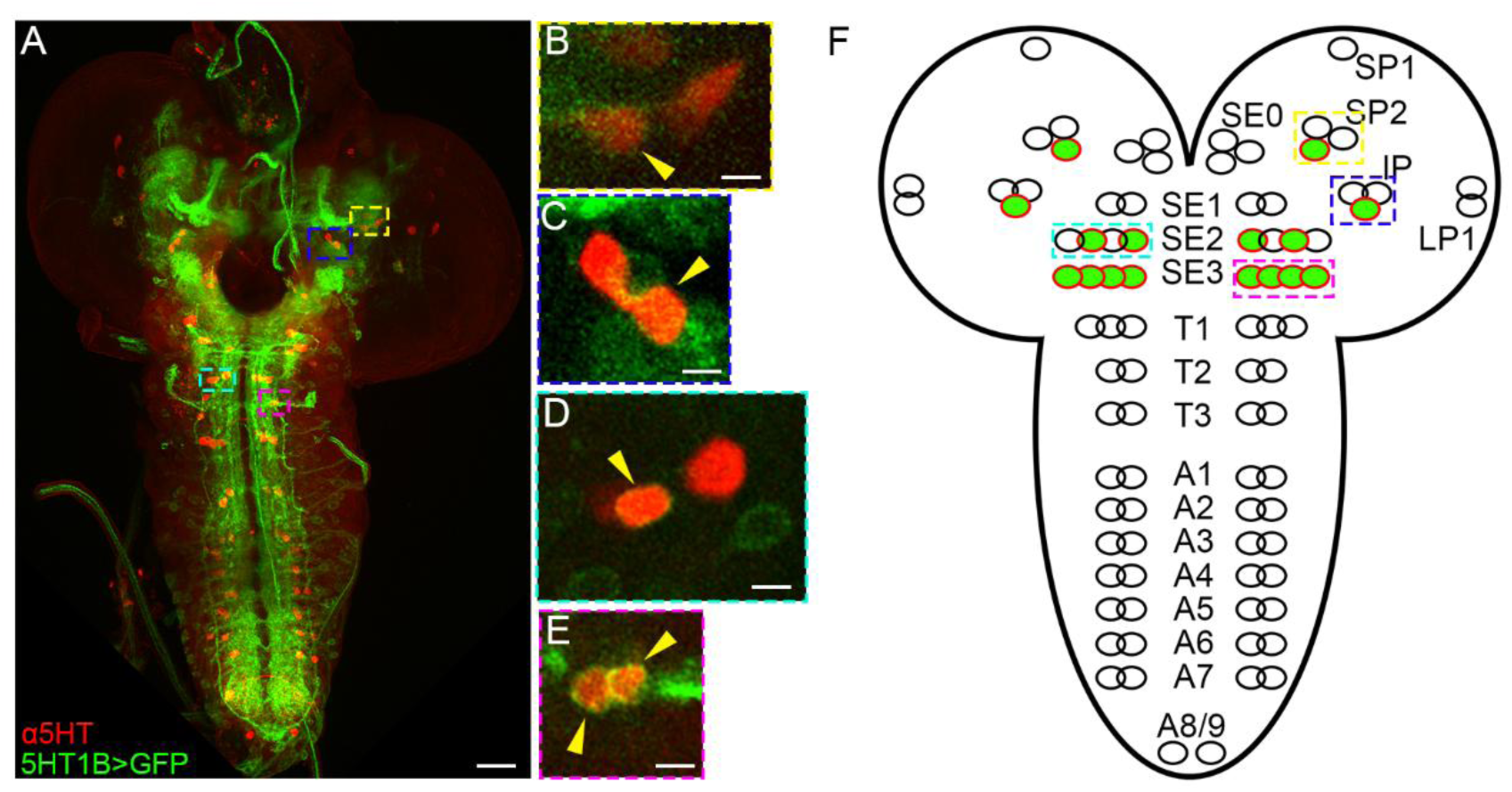
Spatial map of serotonin autoreceptor 5-HT1B expression in third instar brains by genetic labeling. (**A-E**) Images from 5-HT1B-LexA; LexAop-GFP brains stained with ɑ5HT antibodies, with 5-HT1B-expressing serotonergic neurons identified with yellow arrowheads. (**A**) Maximum projection of the Z-stack shows the anatomical layout of the serotonergic system (red) together with neurons that express 5-HT1B (green). (**B**) Magnified view of the SP2 cluster, showing a serotonergic neuron expressing 5-HT1B. (**C**) IP1 serotonergic neuron expressing 5- HT1B. (**D**) SE2 serotonergic neurons expressing 5-HT1B. (**E**) SE3 serotonergic neurons expressing 5-HT1B. (**F**) Schematic of a larval brain with approximated serotonergic clusters. Neurons colorized red and green indicate identified serotonergic neurons expressing 5-HT1B in at least 75% of brains scored. Incompletely colorized clusters indicate some but not all serotonergic neurons express 5-HT1B in that cluster, but do not necessarily represent the specific anatomical position within that cluster. Colored boxes in (A-F) identify the same clusters across panels. Scale bar in (A) = 25 µm; scale bar in (B-E) = 5 µm.

**Figure 4:**
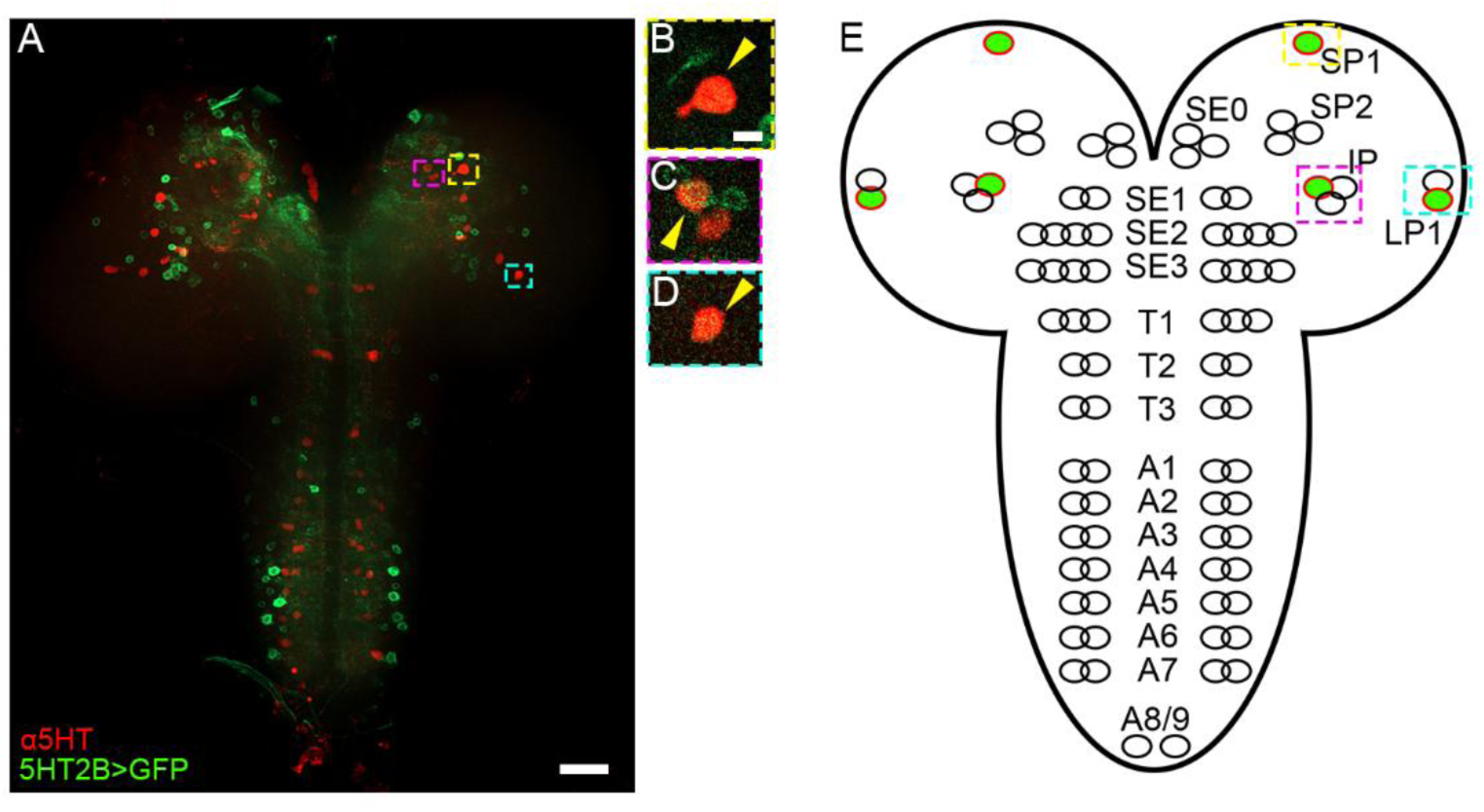
Spatial map of serotonin autoreceptor 5-HT2B expression in third instar brains by genetic labeling. (**A-E**) Images from 5-HT2B-LexA; LexAop-GFP brains stained with ɑ5HT antibodies, with 5-HT2B-expressing serotonergic neurons identified with yellow arrowheads. (**A**) Maximum projection of the Z-stack shows the anatomical layout of the serotonergic system (red) together with neurons that express 5-HT2B (green). (**B**) Magnified view of the SP1 serotonergic neuron expressing 5-HT1B. (**C**) IP1 cluster showing one serotonergic neuron expressing 5-HT2B. (**D**) LP1 serotonergic neuron expressing 5-HT2B. (**E**) Schematic of a larval brain with approximated serotonergic clusters. Neurons colorized red and green indicate identified serotonergic neurons expressing 5-HT2B in at least 75% of brains scored. Incompletely colorized clusters indicate some but not all serotonergic neurons express 5-HT1B in that cluster, but do not necessarily represent the specific anatomical position within that cluster. Colored boxes in (A-E) identify the same clusters across panels. Scale bar in (A) = 25 µm; scale bar in (B-D) = 5 µm.

**Figure 5:**
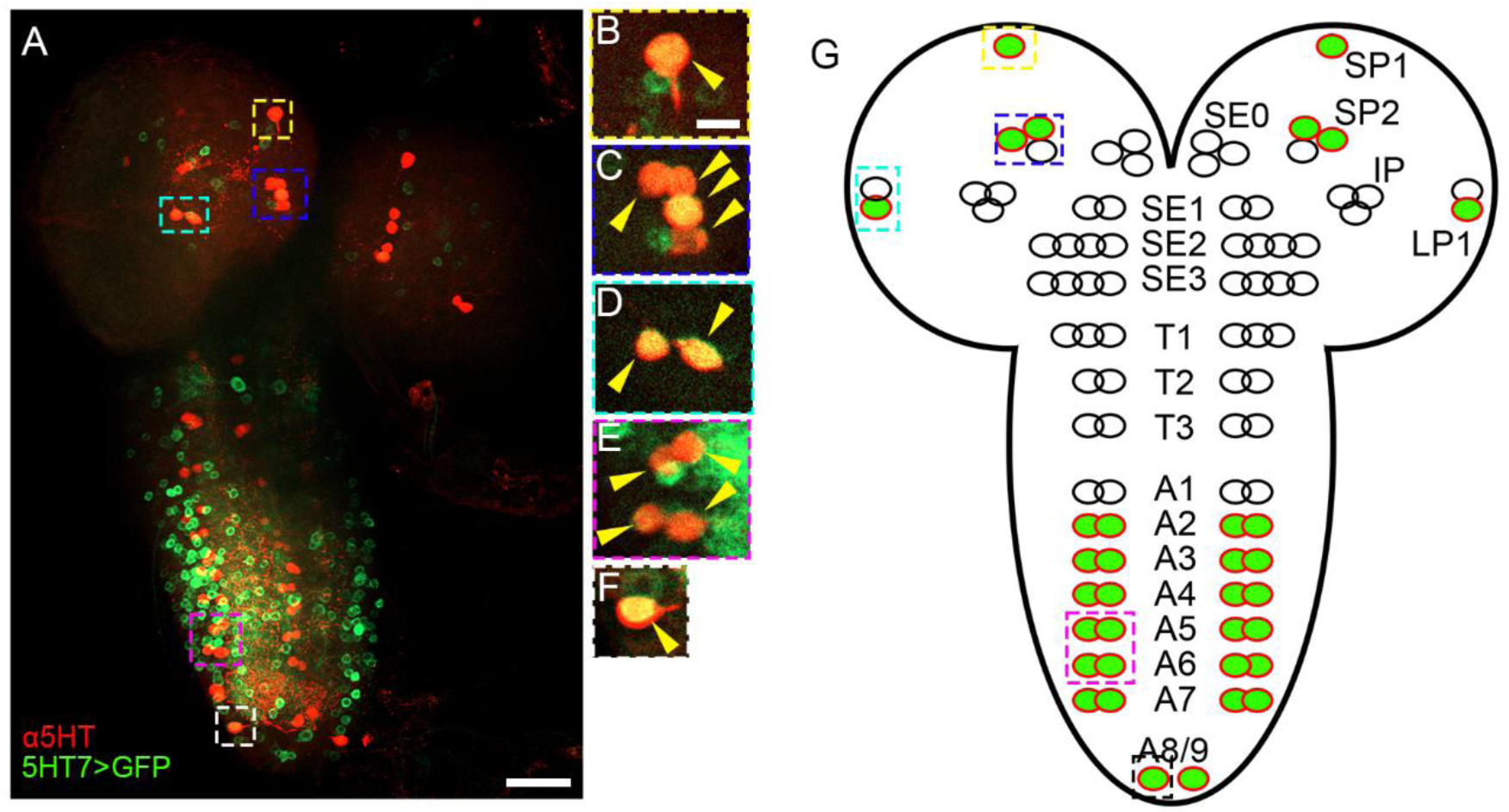
Spatial map of serotonin autoreceptor 5-HT7 expression in third instar brains by genetic labeling. (**A-E**) Images from 5-HT7-LexA; LexAop-GFP brains stained with ɑ5HT antibodies, with 5-HT7-expressing serotonergic neurons identified with yellow arrowheads. (**A**) Maximum projection of the Z-stack shows the anatomical layout of the serotonergic system (red) together with neurons that express 5-HT7 (green). (**B**) Magnified view of the SP1 serotonergic neuron, which expresses 5-HT7. (**C**) SP2 serotonergic neuron cluster expressing 5-HT7. (**D**) LP1 serotonergic neurons expressing 5-HT7. (**E**) A5-A6 serotonergic neurons expressing 5-HT7. (**F**) A8/9 serotonergic neuron expressing 5-HT7. (**G**) Schematic of a larval brain with approximated serotonergic clusters. Neurons colorized red and green indicate identified serotonergic neurons expressing 5-HT7 in at least 75% of brains scored. Incompletely colorized clusters indicate some but not all serotonergic neurons express 5-HT7 in that cluster, but do not necessarily represent the specific anatomical position within that cluster. Note that A2-A8/9 showed 5-HT7 expression though only A5-6 are shown in (E) for simplicity. Colored boxes in (A-F) identify the same clusters across panels. Scale bar in (A) = 25 µm; scale bar in (B-E) = 5 µm.

**Table 3:** Summary of serotonin autoreceptor expression in serotonergic clusters in the larval brain. + indicates one or more serotonergic neurons in that cluster express that specific serotonin receptor. - indicates no serotonergic neurons were discovered in that serotonergic cluster that express that specific serotonin receptor.

|  | SP1 | SP2 | IP | LP1 | SE0-3 | T1-3 | A1-9 |
| --- | --- | --- | --- | --- | --- | --- | --- |
| **1A** | + | + | + | + | - | - | - |
| **1B** | - | + | + | - | + | - | - |
| **2A** | - | - | - | - | - | - | - |
| **2B** | + | - | + | + | - | - | - |
| **7** | + | + | - | + | - | - | + |

Both the larval and adult stages of *Drosophila* metamorphosis are useful models of behavior, with a variety of behaviors modulated by serotonin. Previously, 5-HT1A and 5-HT1B were identified as autoreceptors in adults by genetic labeling (Sampson et al., 2020), prompting us to take a similar approach to identifying serotonin autoreceptors in the adult brain. We accessed and assessed adult single-cell RNA transcriptomic datasets that are publicly available through Fly Cell Atlas (Li et al., 2022) and SCope (https://scope.aertslab.org/), and proceeded with data from Davie, Jannssens and Koldere et al., 2018 for its strong coverage and quality (Davie et al., 2018). From the full dataset of 118,687 cells, we were able to perform *in silico* selection of serotonergic cells using classic markers Trh, SerT, and Vmat. Focused on the 4632 selected cells, we established new nearest-neighbor calculations and Leiden clustering, yielding 16 subclusters with clearly bordered UMAP representations (**Fig 6A**). The five receptors showed obvious differential expression, where 5-HT1A showed the broadest coverage in 31.6% of the serotonergic neurons, followed by 5-HT2A and 5-HT1B at 12.9%, 12.5%, respectively, and 5-HT7, 5-HT2B’s expression percentages were much lower, at 9.4%, 7.0%, respectively. Our refined clustering analysis also hinted the potential of receptor co-expression (e.g., clusters 2, 11 and 13), however, the actual observable co-expression in real biological samples could be very limited due to the highly variable expression levels of these receptors (**Fig 6B**).

**Figure 6:**
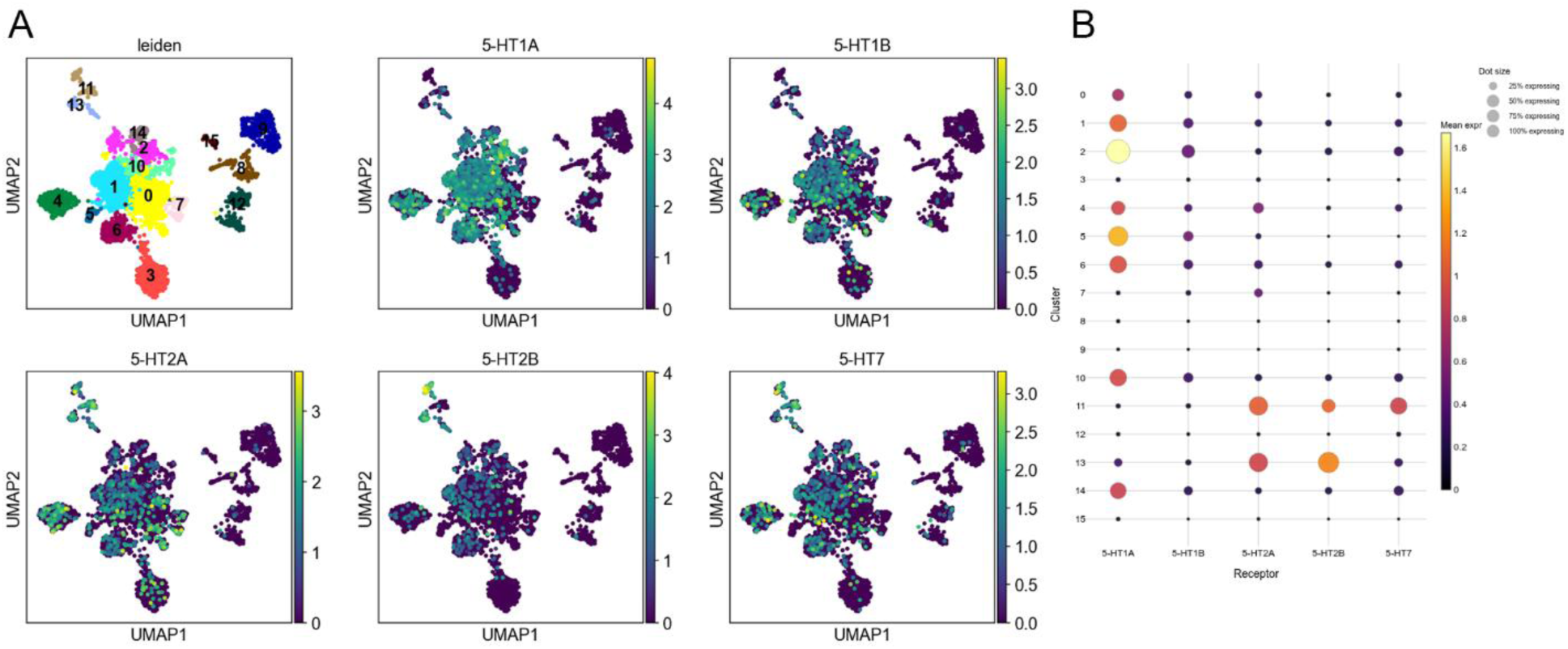
Analysis of single-cell sequencing data from adult heads shows all five serotonin receptors are expressed in serotonergic neurons. (**A**) Leiden cluster identity of adult serotonergic neurons, filtered and re-clustered *in silico* from the full dataset of Davie et al 2018, as well as the expression distribution patterns of all five serotonin receptors, are displayed with a UMAP embedding in 2D. (**B**) Per-cluster expression profile of 5-HT receptors. The mean expression level of a given receptor in a given cluster is displayed via color depth in each dot, showcased by the color bar. The proportion of cell population in a given cluster expressing a given receptor is displayed through the size of each dot.

We used the same genetic labeling strategy to determine which serotonergic neurons in the adult brain express serotonin receptors. In the adult central brain, we found 5-HT1A expressed in PMPM (**Fig 7B**), LP (**Fig 7C**), and both anterior and posterior SEM clusters (**Fig 7D-E**); 5-HT1B was expressed in SEL (**Fig 8B**), AMP (**Fig 8C**), and LP (**Fig 8D**); 5-HT2B was expressed in PMPD (**Fig 9B**) and PMPM (**Fig 9C**); and 5-HT7 was only expressed in LP serotonergic neurons (**Fig 10**). As with the larval brains, we were not able to find GFP positive serotonergic neurons in adult brains. These data are summarized in Table 4.

**Figure 7:**
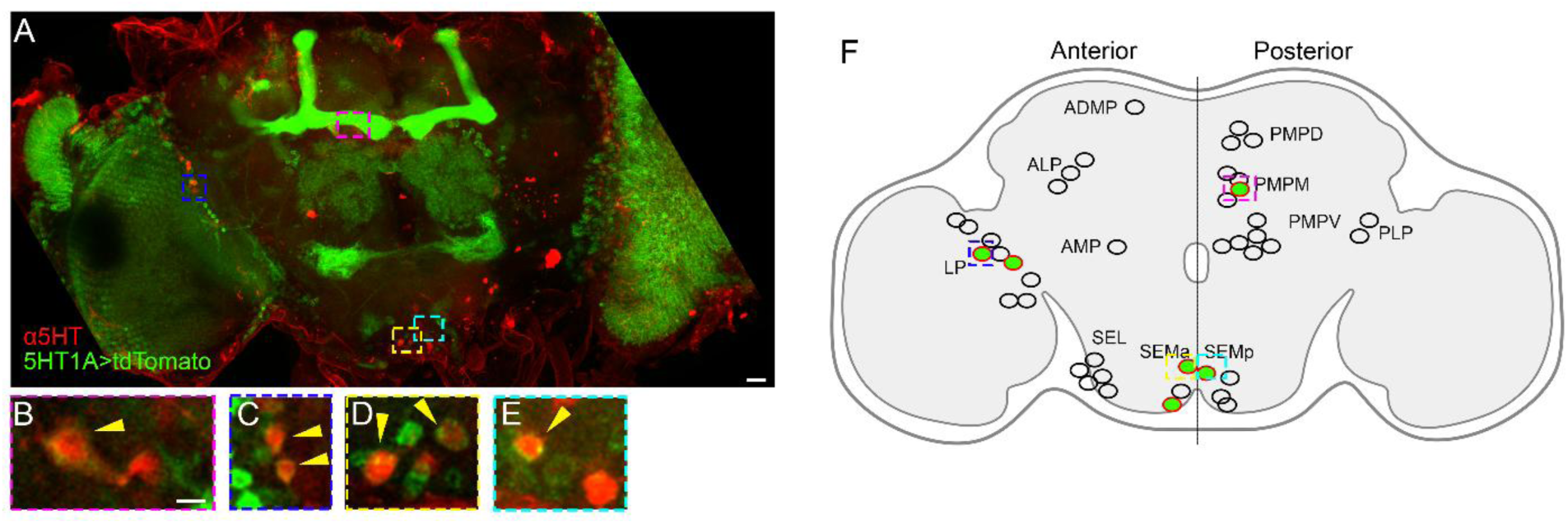
Spatial map of serotonin autoreceptor 5-HT1A expression in adult brains by genetic labeling. (**A-F**) Images from 5-HT1A-LexA; LexAop-GFP brains stained with ɑ5HT antibodies, with 5-HT1A-expressing serotonergic neurons identified with yellow arrowheads. (**A**) Maximum projection of the Z-stack shows the anatomical layout of the serotonergic system (red) together with neurons that express 5-HT1A (green). (**B**) Magnified view of the PMPM cluster, showing a serotonergic neuron expressing 5-HT1A. (**C**) LP1 serotonergic neurons expressing 5- HT1A. (**D**) SEMa serotonergic neurons expressing 5-HT1A. (**E**) SEMp serotonergic neurons expressing 5-HT1A. (**F**) Schematic of an adult brain with approximated serotonergic clusters. Neurons colorized red and green indicate identified serotonergic neurons expressing 5-HT1A in at least 75% of brains scored. Incompletely colorized clusters indicate some but not all serotonergic neurons express 5-HT1A in that cluster, but do not necessarily represent the specific anatomical position within that cluster. Colored boxes in (A-F) identify the same clusters across panels. Scale bar in (A) = 25 µm; scale bar in (B-E) = 5 µm.

**Figure 8:**
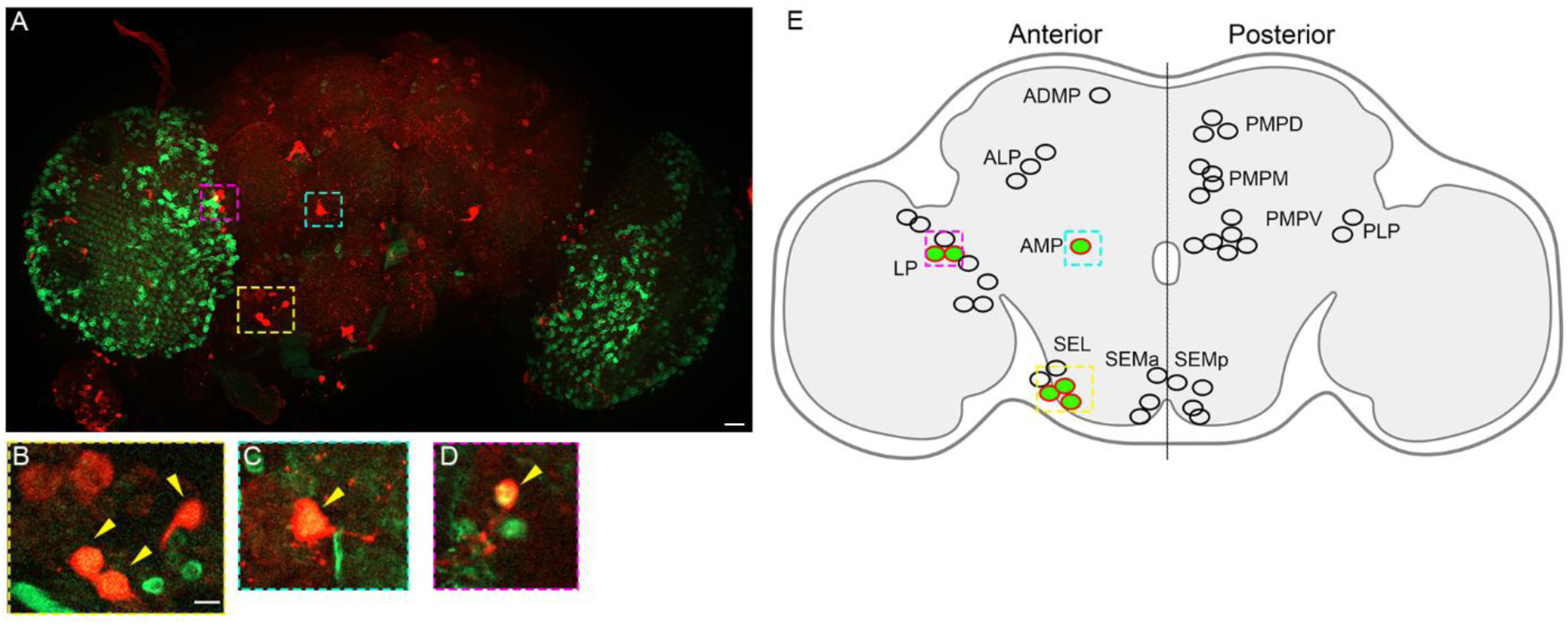
Spatial map of serotonin autoreceptor 5-HT1B expression in adult brains by genetic labeling. (**A-F**) Images from 5-HT1B-LexA; LexAop-GFP brains stained with ɑ5HT antibodies, with 5-HT1B-expressing serotonergic neurons identified with yellow arrowheads. (**A**) Maximum projection of the Z-stack shows the anatomical layout of the serotonergic system (red) together with neurons that express 5-HT1B (green). (**B**) Magnified view of the SEL cluster, showing a serotonergic neuron expressing 5-HT1B. (**C**) AMP serotonergic neuron expressing 5- HT1B. (**D**) LP serotonergic neuron expressing 5-HT1B. (**E**) Schematic of an adult brain with approximated serotonergic clusters. Neurons colorized red and green indicate identified serotonergic neurons expressing 5-HT1B in at least 75% of brains scored. Incompletely colorized clusters indicate some but not all serotonergic neurons express 5-HT1B in that cluster, but do not necessarily represent the specific anatomical position within that cluster. Colored boxes in (A-E) identify the same clusters across panels. Scale bar in (A) = 25 µm; scale bar in (B-D) = 5 µm.

**Figure 9:**
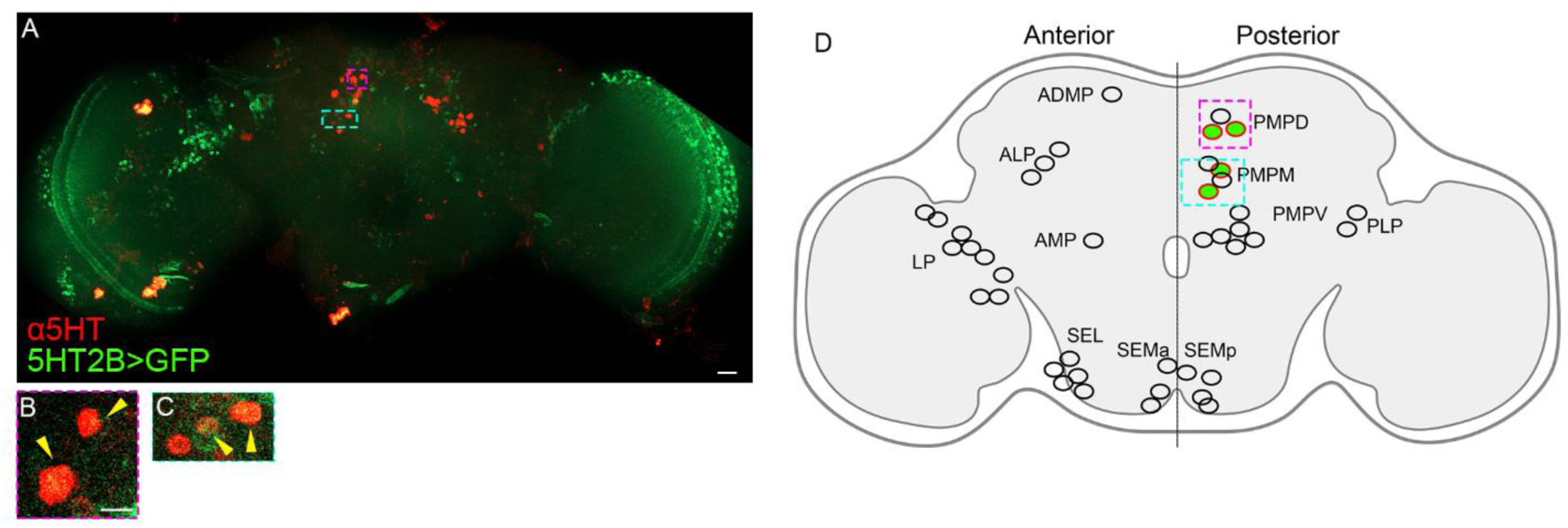
Spatial map of serotonin autoreceptor 5-HT2B expression in adult brains by genetic labeling. (**A-F**) Images from 5-HT2B-LexA; LexAop-GFP brains stained with ɑ5HT antibodies, with 5-HT1B-expressing serotonergic neurons identified with yellow arrowheads. (**A**) Maximum projection of the Z-stack shows the anatomical layout of the serotonergic system (red) together with neurons that express 5-HT2B (green). (**B**) PMPD cluster, showing serotonergic neurons expressing 5-HT2B. (**C**) PMPM cluster, showing serotonergic neurons expressing 5- HT2B. (**D**) Schematic of an adult brain with approximated serotonergic clusters. Neurons colorized red and green indicate identified serotonergic neurons expressing 5-HT2B in at least 75% of brains scored. Incompletely colorized clusters indicate some but not all serotonergic neurons express 5-HT2B in that cluster, but do not necessarily represent the specific anatomical position within that cluster. Colored boxes in (A-D) identify the same clusters across panels. Scale bar in (A) = 25 µm; scale bar in (B-C) = 5 µm.

**Figure 10:**
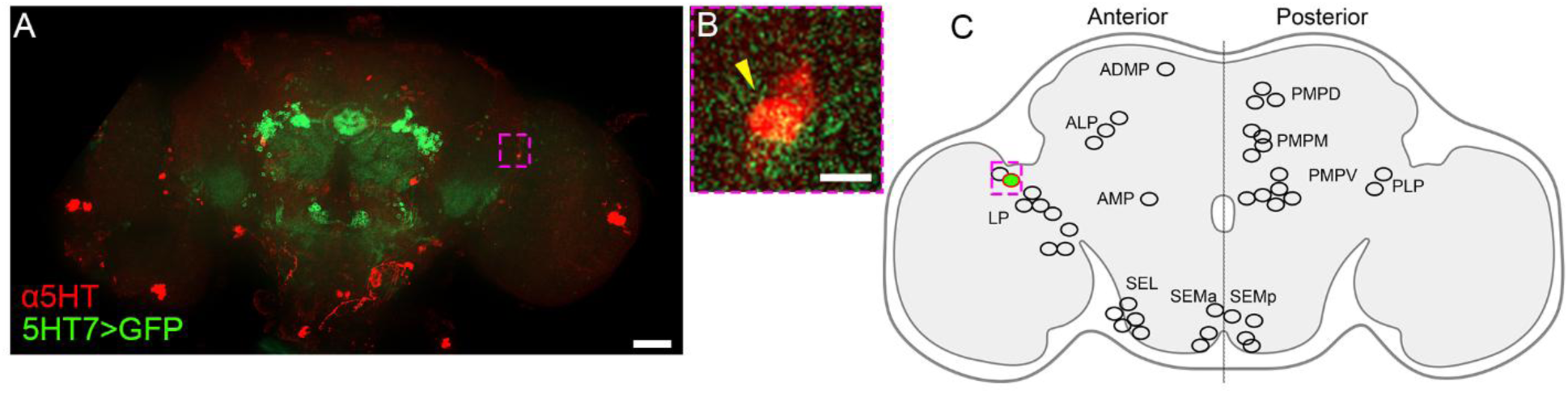
Spatial map of serotonin autoreceptor 5-HT7 expression in adult brains by genetic labeling. (**A-C**) Images from 5-HT7-LexA; LexAop-GFP brains stained with ɑ5HT antibodies, with 5-HT7-expressing serotonergic neurons identified with yellow arrowheads. (**A**) Maximum projection of the Z-stack shows the anatomical layout of the serotonergic system (red) together with neurons that express 5-HT7 (green). (**B**) Magnified view of the LP cluster, showing a serotonergic neuron expressing 5-HT7. (**C**) Schematic of an adult brain with approximated serotonergic clusters. Neurons colorized red and green indicate identified serotonergic neurons expressing 5-HT7 in at least 75% of brains scored. Incompletely colorized clusters indicate some but not all serotonergic neurons express 5-HT7 in that cluster, but do not necessarily represent the specific anatomical position within that cluster. Colored boxes in (A-C) identify the same clusters across panels. Scale bar in (A) = 25 µm; scale bar in (B-D) = 5 µm.

**Table 4:** Summary of serotonin autoreceptor expression in serotonergic clusters in the adult brain. + indicates one or more serotonergic neurons in that cluster express that specific serotonin receptor. - indicates no serotonergic neurons were discovered in that serotonergic cluster that express that specific serotonin receptor. For simplicity, only clusters with identified positive hits are included in this table.

|  | LP | AMP | SEMa | SEL | PMPD | PMPM | SEMp |
| --- | --- | --- | --- | --- | --- | --- | --- |
| 1A | + | - | + | - | - | + | + |
| 1B | + | + | - | + | - | - | - |
| 2A | - | - | - | - | - | - | - |
| 2B | - | - | - | - | + | + | - |
| 7 | + | - | - | - | - | - | - |

## Discussion

Here we provide evidence that all five of the serotonin receptors in the *Drosophila* genome are expressed in serotonergic neurons (autoreceptors). This has been demonstrated in two ways: our sequencing results at both the larval and adult stages; and our genetic labeling results showing four of the five serotonin receptors expressed, with the exception being 5-HT2A. Structurally, the five serotonin receptor genes in *Drosophila* are assumed to be orthologs of mammalian 5-HT1A, 5-HT1B, 5-HT2, and 5-HT7 (Blenau et al., 2017; Colas et al., 1995, 1997; Kasture et al., 2018; Saudou et al., 1992). While we found all five of serotonin receptor genes are expressed in serotonergic neurons, there are some discrepancies between this and what has been found in mammalian systems. Namely, 5-HT7 has not been identified as an autoreceptor in mammals. Conversely, 5-HT5A is expressed in brainstem neurons in chicken (Fujita et al., 2022), though this is not believed to have an orthologous gene in *Drosophila*. Serotonin receptors share overlapping signaling pathways between invertebrates and mammals as well: *Drosophila* 5-HT1A stimulates the opening of G-protein-coupled inward rectifying potassium channels to inhibit serotonin secretion (Witz et al., 1990); *Drosophila* 5-HT1B couples to Gi/Go G-Proteins to inhibit cAMP production as in mammals (Kasture et al., 2018); activation of *Drosophila* 5-HT2A and 5- HT2B lead to calcium signaling in an inositol-1,4,5-triphosphate dependent manner (Blenau et al., 2017) similar to mammalian 5-HT2; and *Drosophila* 5-HT7 couples to Gs when activated to stimulate cAMP production (Becnel et al., 2011; Blenau et al., 2017).

In general, serotonin initiates signaling pathways that autoregulate the development of serotonergic neurons (Budnik et al., 1989; Daubert & Condron, 2010; Diefenbach et al., 1995; Kinser et al., 2026; Long et al., 2023; Richardson-Jones et al., 2011; Sykes & Condron, 2005) and serotonin secretion in developed circuits (reviewed in Trueta and Cercos 2026). In mammals, serotonergic tone is thought to result from opposing Gi-coupled 5-HT1A/B inhibitory activity and Gq-coupled excitatory activity (McDevitt & Neumaier, 2011; Quentin et al., 2018). Beyond that, much of what we believe about serotonin receptor signaling is derived from experiments using heterologous receptor expression in non-serotonergic or even non-neuronal cells (Rojas & Fiedler, 2016), and there is some evidence that serotonin autoreceptors couple to different G- proteins than their heteroreceptor counterparts (Albert et al., 1999; Penington et al., 1993). A distinct advantage of using *Drosophila* as a model organism to study serotonin autoreceptor signaling is the accessibility of isolated serotonergic neurons with known and variable receptor expression patterns. Not only are serotonergic neurons reasonably isolated *in vivo* compared to the dense raphe nucleus, but primary serotonergic neurons can be grown and visually isolated in culture (Long et al., 2023). Moreover, knowing the specific identity of serotonin receptor- expressing serotonergic neurons provide behaviors that serve as putative functional readouts of serotonergic function in mutants. For example, we found serotonergic neurons in LP1 express 5- HT1A, 2B, and 7, and these neurons innervate the larval optic neuropil (Huser et al., 2012; Larderet et al., 2017). Therefore, vision-based behaviors such as phototaxis (Rodriguez Moncalvo & Campos, 2009) or visual attraction (Dombrovski et al., 2019; Justice et al., 2012) can be used in future experiments to determine how mutations in signaling pathways downstream of these receptors affect the function of serotonergic tone.

Our anatomical maps of serotonin autoreceptor expression suggest that there are combinatorial autoreceptor expressions in a variety of serotonergic neuron clusters throughout the *Drosophila* nervous system, though our approach precludes us making this as a definitive conclusion for all clusters. The SP1 neuron was positive for 5-HT1A, 5-HT2B, and 5-HT7 and it is the only serotonergic neuron in that anatomical cluster. It is therefore reasonable to conclude that these three autoreceptors are co-expressed in SP1. However, we used genetic labeling of a single receptor subtype expression per brain, and we identified positive clusters as containing one or more double-positive serotonergic neurons and did not delineate between serotonergic neurons within the cluster. Therefore, we can not distinguish between neurons co-expressing autoreceptors and expression in different neurons within the cluster. Future experiments using combinatorial labeling to provide direct evidence and anatomical information for autoreceptor co- expression.

Our genetic labeling approach was advantageous for this study by allowing us to visualize fluorescent cell bodies. Cell bodies are large discrete objects that are easily identifiable and registered to anatomical maps compared to individual receptor proteins. However, there are two notable limitations. First, we are unable to localize serotonin autoreceptors within serotonergic neurons. This is not trivial, as serotonin has different effects on leech neurons depending on the neuron compartment it acts on due to varying localizations of serotonin autoreceptors (Cercós et al., 2009; Leon-Pinzon et al., 2014; Trueta et al., 2004; Trueta & Cercós, 2026). Second, we found a lower representation of serotonergic neurons expressing autoreceptors in our spatial experiments compared to what was expected from the sequencing results. This is obvious for 5- HT2A, where we were unable to find double-positive serotonergic neurons. While we cannot rule out the possibility that 5-HT2A is not an autoreceptor in *Drosophila*, we believe this discrepancy is due to the low sensitivity of genetic labeling rather than false-positives in both mRNA sequencing datasets. This is likely also the reason we did not find 5-HT1A-expressing serotonergic neurons in SEL or PMP clusters, nor 5-HT1B in the PMP neurons as was shown previously (Sampson et al., 2020). We believe that these discrepancies are a result of different genetic labeling systems, Gal4 versus LexA, and therefore do not preclude these as being autoreceptor expressing serotonergic neurons. Since our approach errs on the side of false negatives, we feel confident that these maps can be used to generate new hypotheses for autoreceptor function, though perhaps not comprehensively.

